# Fabrication and Evaluation of Mechanical and Tribological Properties of ultra-high-molecular-weight polyethylene-based Nanocomposites Reinforced with Hydroxyapatite, Zirconia and Multi-walled Carbon Nanotubes

**DOI:** 10.1101/2021.05.03.442451

**Authors:** Alireza Nikbakht, Jafar Javadpour, M. Reza Naimi-Jamal, Hamidreza Rezaie

## Abstract

The unique properties of ultra-high-molecular-weight polyethylene (UHMWPE) have made composites based on this polymer the gold standard choice for articulating surfaces used in arthroplasty. However, wear debris released by prosthesis is still a major concern of the implants. In this study, we address the urgent need to revisit the current methodologies used in designing these biomaterials by fabricating UHMWPE-based nanocomposite with Hydroxyapatite (HA), Multi-walled Carbon Nanotubes (MWCNTs), and Zirconia as additives. We investigated how different combinations of these additives impact the nanocomposites’ hardness, plasticity index (PI), and friction coefficient. Our results show a constant increase in hardness by increasing Zirconia. The MTT assay test and Scanning Electron Microscopy (SEM) demonstrated an increase in cell attachment and cell viability. Examining various additives’ amounts, we can further explore the possibility of reaching an optimum proportion of the ingredients. Compared with other UHMWPE-based nanocomposites (UHMWPE-HA-MWCNT’s), the fabricated nanocomposite shows an improvement in tribological properties.

## 1. Introduction

An ageing population, and the growing number of age-associated diseases such as osteoarthritis, osteoporosis, and physical injuries worldwide has led to the rise in joint replacement surgeries, particularly in the lower limbs. Joint replacement surgeries may cause pain relief, improve walking ability, and boost patients’ morale. However, the negative effect of wear debris of implants’ bearing surfaces in total joint replacement and consequent inflammation resulting from Ultra-high-molecular-weight polyethylene (UHMWPE) byproducts has always been a critical issue for researchers and patients [**1**].

American Academy of Orthopaedic Surgeons (AAOS) reported 142,000 total hip replacements, 108,000 partial hip replacements, and 418,000 total knee replacements in 2007 in the United States alone. It is also projected that the number of people undergoing knee and hip replacement surgeries every year will surge to 3.48 million and 572,000, respectively, by 2030 [**2**].

Choosing the ideal materials and finding the optimum processing method to design an appropriate prosthesis is one of the pressing issues in arthroplasty applications in recent years. So, there are numerous investigations of biomaterials and their suitability for prosthetics. For example, bioceramic composites with HA matrix and a wide range of novel additives like MWCNTs were widely popular just a few years ago [**3**].

Among the most used biomaterials, UHMWPE is one of the best-known materials used as cups in total hip replacements, owing to its biocompatibility and desirable tribological behaviour [**4**].

Consequently, numerous studies on this polymer have been revealed to improve its tribological and mechanical properties and address some of its limitations in joint replacement surgeries [**5**].

Several studies on using gamma irradiation [**6**], applying different processing methods [**7,8**], adding vitamin E [**6**,**9**], and changes in crystallinity have been carried out to improve wear resistance and mechanical properties (through increasing cross-linking and preventing oxidation) of this well-known polymer.

Fabricating UHMWPE-based composite, using different types of additives such as HA [**9**], Carbon Nanofibers [**10**], Carbon Nanotubes [**11**], Zirconia [**12,13**], and Alumina [**14**] or even using some additives such as Zirconia and HA at the same time [**15**] in order to reduce the wear rate of this polymer has been widely investigated. Recent evidence suggests that the powder mixing method and the fabrication method may significantly impact the final composites’ properties [**10**,**16**]. On the other hand, harvesting UHMWPE-based composite’s full capacity in orthopaedic applications culminated in drawing researchers’ attention to fabricating hybrid composite with a wide range of manageable properties [**17**]. This view is supported by Sattari et al. [**18**]. They reported an improvement in hardness and Young’s modulus in UHMWPE-SCF (short carbon fibres)/nano-SiO_2_ (silica) hybrid composites.

Mechanical properties and hardness were also investigated via nanoindentation analysis by Gupta et al. in 2013 for compression moulded HA-MWCNTs-Al_2_O_3_-UHMWPE hybrid composites, and the results demonstrated improvements of mechanical properties by 3%-5% with the Al_2_O_3_ addition. Weak polymer additives’ interface (HA and CNT), followed by a decline in mechanical properties due to a lack of a coupling agent were shown. In contrast, in samples at sub-micron size, addition of Al_2_O_3_ powder as a densifying factor depicted improved mechanical properties [**14**].

Taking into account previous studies on enhanced UHMWPE properties, and its applications in arthroplasty, reinforced HA-MWCNTs-ZrO_2_-UHMWPE nanocomposites were fabricated to investigate the effect of ZrO_2_ on the tribological and mechanical behaviour of designed nanocomposites. Also, the nanocomposites’ cytocompatibility was proved by both MTT assay and cell adhesion analysis.

## 2. Experimental Materials

UHMWPE powder with the mean particle size of 150 microns, the average molecular weight between 3,000,000 and 6,000,000, and density of 94 *g/cm*^3^ (in 25 centigrade degree), and Hexane with the purity of 99% were purchased from Merck. Calcium nitrate tetrahydrate and di-ammonium hydrogen phosphate were used as a source of calcium and phosphate, respectively. 25% ammonia solution was also supplied from Merck to use in HA synthesis. Multi-Walled Carbon Nanotubes with a purity of over 95% (outer diameter of 20-30 nm, the inner diameter of 5-10 nm, length up to 30 nm, and an ash content of less than 1.5%.) were purchased from the US Research Nanomaterials, Inc. for further functionalization.

Zirconia powder with a purity of 99.5% and a particle size of 5 μm was purchased from the Western Australia company. Liquid Paraffin Pharma-Grade was purchased from Raha Paraffine Company to use for nanocomposites fabrication in the melt mixing process.

Roswell Park Memorial Institute (RPMI) 1640 Medium, Fetal Bovine Serum (FBS), and Trypsin-EDTA solution were supplied from GIBCO. Methylthiazolyldiphenyl-tetrazolium bromide (MTT), Dimethyl sulfoxide (DMSO), Phosphate Buffered Saline (PBS), and Glutaraldehyde were purchased from Sigma Aldrich. Bone cell line MG-63 was provided by the Pasteur Institute of Iran’s Cell Bank.

## 3. Experimental

### 3.1. HA synthesis

Nano-sized HA powder was synthesized through the wet-chemical precipitation process based on HA synthesis’s Stoichiometric reaction (1). The pH was adjusted to 10 with *N H*4*OH* solution 25%, and the Ca/P ratio was observed at 1.67 by stoichiometry [**19**].

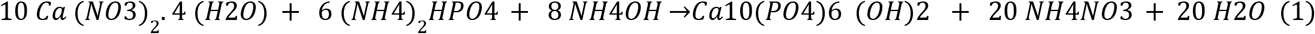

A diluted solution was obtained from mixing two salt solutions containing diammonium hydrogen phosphate and calcium nitrate tetrahydrate nitrate, stirred at 40 °C for one hour at 400 rpm. The resulting solution then was aged for 24 hours at room temperature.

Final milky solutions were heated up to 800 °C with a constant heating rate of 10 °C/min.

### 3.2. Oxidation of MWCNT

One gram of MWCNT was initially sonicated in a conventional ultrasonic bath (FRITSCH LABORETTE 17 100 W, 42 kHz) for 30 minutes at room temperature. The solution of 50 ml of nitric acid 65% and 150 ml of sulfuric acid 94% (both acids were supplied from Merck) was then stirred at 60 °C for two hours. Then filtered and washed till the pH of extracted water reached 7. In the end, the purified carbon nanotubes were dried in the oven (BM120E FG Iran) at 80°C overnight. The purpose of this procedure is to help oxidize the surface of materials and improve CNTs interaction and dispersion.

### 3.3. Zirconia size reduction

The Zirconia powder with a micrometre size range underwent the wet-grinding process in the planetary ball mill (Sanat Ceram Mehr Alborz PLM) to reduce Zirconia’s particle size. After passing through a filter with a desired nanometer-size mesh, the nano powder was dried in the oven at 70°C for 24 hours [**20**] and finally prepared to be measured by SEM.

### 3.4. Composite fabrication process

Regarding the solution’s low density, during the first step, the ultrasonic bath was used to mix composite components through sonication, then offering proper dispersion of additives in the composite matrix. The Internal mixing method was used to fabricate the nanocomposites. Composite contents are shown in TABLE1.

**TABLE1:** Composites contents

| Contents |  |  |  |
| --- | --- | --- | --- |
| Sample | ZrO<sub>2</sub> (phr) | CNT (functionalized)(phr) | HA(phr) |
| S0 | 0 | 5/0 | 30 |
| S1 | 3 | 5/0 | 30 |
| S2 | 6 | 5/0 | 30 |
| S3 | 9 | 5/0 | 30 |

The preparation of the composite powder for each sample was performed as follows. First, the functionalized carbon nanotubes with an appropriate amount of ethanol-95% were sonicated in an ultrasonic bath for 15 min to distribute the carbon nanotubes well into ethanol and provide a homogeneous distribution in other parts of the composite. Synthesized HA was then added to the CNT solution and sonicated for further 15 minutes to obtain a homogeneous distribution of additives in the nanocomposites. Paraffin oil was added (15 phr of the composite’s total weight) to decrease the polymer’s viscosity and enhance the additive dispersion. In the last stage, supplied UHMWPE was added when composite solutions were sonicated for another 15 minutes.

All samples were heated at 80 °C for 48 hours for ethanol evaporation. This heat and time help paraffin molecules penetrate the polymer particles (all four composite specimens were treated in the same way). Following this, the initial composite powder was ready for the melt mixing process.

The two-roll mill internal mixer was used to blend the filler particles homogeneously with adding paraffin oil (15 phr of the composite) for swelling of UHMWPE. Then the nanopowder composite constituents were mixed at 210 °C for 2 hours with 150 rpm roll mill speed. The viscosity reduction of the pre-prepared composite nanopowder was achieved during this stage. Then all samples were cooled at room temperature.

All these steps were performed equally for all composite samples. The samples were pressed at 130 °C and an initial pressure of 4 MPa with a pressure increasing rate of 0.5 MPa per minute for 30 minutes to remove paraffin oil from the composites. Each sample was subjected to this hot press process four times. The lower melting temperature of paraffin and the applied pressure led to most paraffin oil being removed from the composite at this stage. After that, the ultra-centrifugal mill ZM200 was used to reduce the particle size of pressed composite bulks to under 100 microns. Reduced size composite powder was stirred at 60 °C with hexane, followed by drying at 80 °C for 24 hrs, to eliminate paraffine.

Following these steps, cylindrical-shaped nanocomposite fabricated through compression molding by atmospheric hot press machine (KOVACO Hotpress KHP-2000 at 210 °C under 10 MPa pressure with a constant heating rate of 5 °C /min) for Nano-indentation and Nanoscratch tests following ISO 14577 and ISO 10993- 5 for cytotoxicity and cell adhesion tests (tablets with 8 mm in diameter and less than 2 cm in thickness).

## 4. Calculation

The samples were polished and sandpapered before carrying out Nanoindentation and Nano Scratch tests. Nanoindentation tests were carried out using a *TriboScope*^®^ system (Hysitron, USA) equipped with a Berkovich type indenter tip in accordance with the procedure described in ISO14577.

### 4.1. Nanoindentation test

Nanoindentation test based on ISO14577 is regarded as one of the novel tests in the mechanical properties investigation. According to Oliver and Pharr’s method [**21**]:

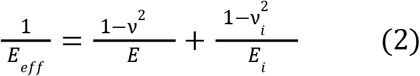

The values of *E_i_*, (elastic modulus) and *v_i_*, (Poisson ratio of the diamond indenter) are 1140 GPa and 0.07, respectively, in eq. 2 [**22**].

Therefore, compressive elastic modulus can be calculated by substituting the value of v, which is reported at the level of 0.4 for pure UHMWPE [**23**].

As the indenter’s stiffness is much greater than that of the composite sample, the sample’s hardness can be calculated as follows [**22**]:

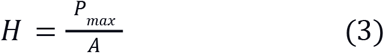

In this equation, *H* is the hardness, and *P_max_* is the maximum force. *max*

### 4.2. Nanoscratch Tests

This test was conducted through the nanoindentation technique, determining the relative material’s resistance to scratch, and investigating the nanocomposites’ tribological properties.

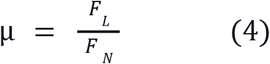

Vertical and lateral force (*F_N_, F_L_*) and displacement were recorded throughout the nanoindentation test. Previous studies have reported that the ratio of lateral forces to vertical forces presenting the coefficient of friction [**22**].

### 4.3. Plasticity index

The plasticity index (PI) is a parameter used to evaluate the elastic-plastic behaviour of materials. The plasticity index correlated to S1 and S2 representing the total indentation work and elastic deformation work, respectively. Plasticity Index can be achieved using the equation from [**22**].

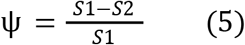

### 4.4. Cytocompatibility test

#### 4.4.1. Sample preparation

The cylindrical specimens were prepared through the hot press’s compression molding process (8 mm in diameter and less than 1 cm in thickness), then polished and sterilized with 70% ethanol and UV exposure for 24 hours.

#### 4.4.2. MTT assay

This test is based on the ability of the mitochondrial dehydrogenase enzymes in living cells to convert the water-soluble yellow dimethylthiazol diphenyltetrazolium bromide to insoluble violet formazan (ISO 10993-5). The absorbance was measured at 570 nm by Elisa Reader (Powerwave xs - BioTeK), and cell viability was compared with an extracted-free well containing 10000 Mg63 cell lines. The procedure was repeated three times for each sample.

#### 4.4.3. Cell adhesion assay

The produced nanocomposite specimens were cut in half to fit a 96-well plate to investigate cell adhesion. Culture medium and approximately seventy thousand cells were added to all four samples.

## 6. Results and Discussion

### 6.1. HA Particles Size Assessment

The synthesized HA particles were subjected to X-ray diffraction for structural evaluation. There are very sharp and high-intensity peaks that indicate the high crystallinity of single-phase HA as shown in the X-ray diffraction pattern of the synthesized HA particles (Figure 1). This test’s results were analyzed, plotted, and checked by Plus Highscore, Anaconda software, and checked with the JCPDS reference card. Despite the matching synthesized HA diffraction pattern with the reference card, there is likely to be a small amount of calcium phosphate and beta-tricalcium phosphate, which are very small and would not affect our material properties.

**Fig. 1.**
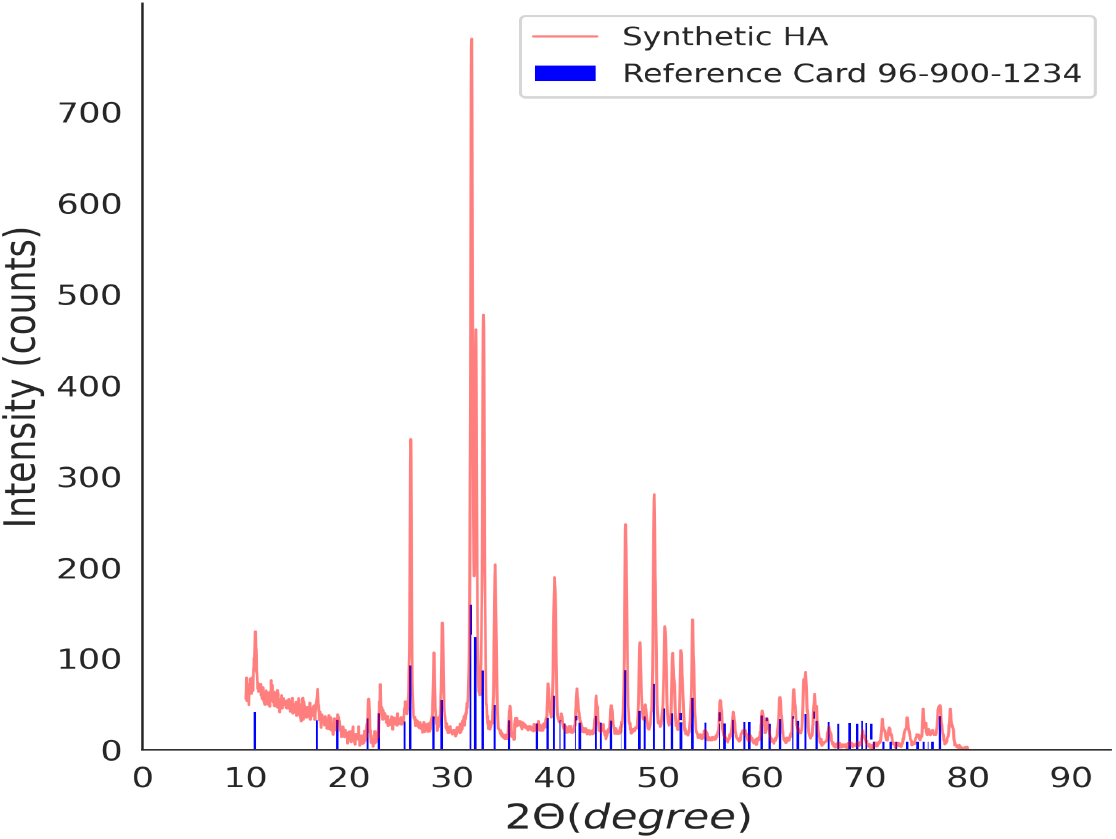
X-ray diffraction pattern of the synthesized HA particles through a Wet-Chemical Precipitation process (red line) in comparison with the JCPD HA reference card (blue line).

The average crystallite size was also measured by using the Scherrer equation and utilizing HighScore plus software[**24**]. The average crystallite size at 6 peaks with maximum intensity revealed to be 27 nm.

As Figure 2 clarifies, the average size of synthesized HA particles through the wet chemical precipitation method is in the nanometer range.

**Fig. 2.**
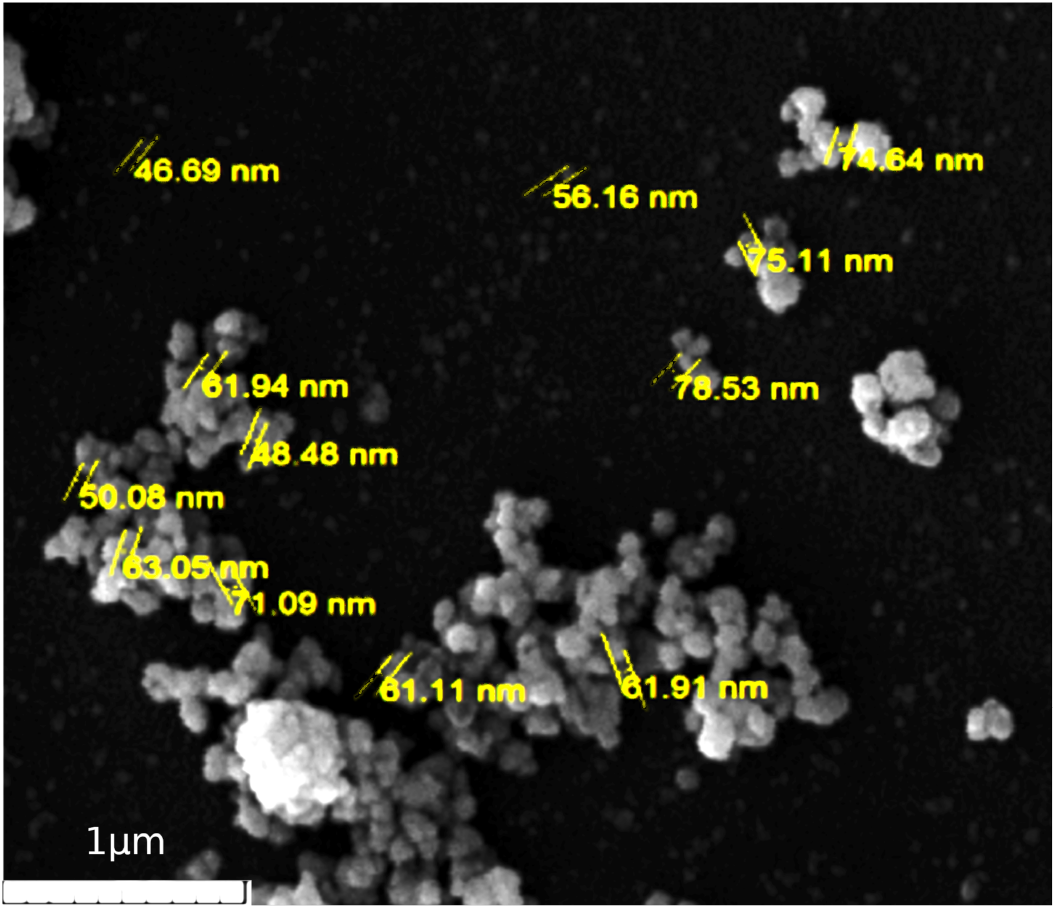
Scanning Electron Microscope image of synthesized HA particles by Wet-Chemical Precipitation process (Avg. Size 60nm, 60,000 x magnification).

### 6.2. Zirconia Particle Size Assessment

As shown in Figure 3, the Zirconia particles were reduced to sub-micron size by approximately 100 nm on average, using the wet grinding method. It should be noted, however, that the smaller the particle size, the greater the tendency of the particles to agglomerate.

**Fig. 3.**
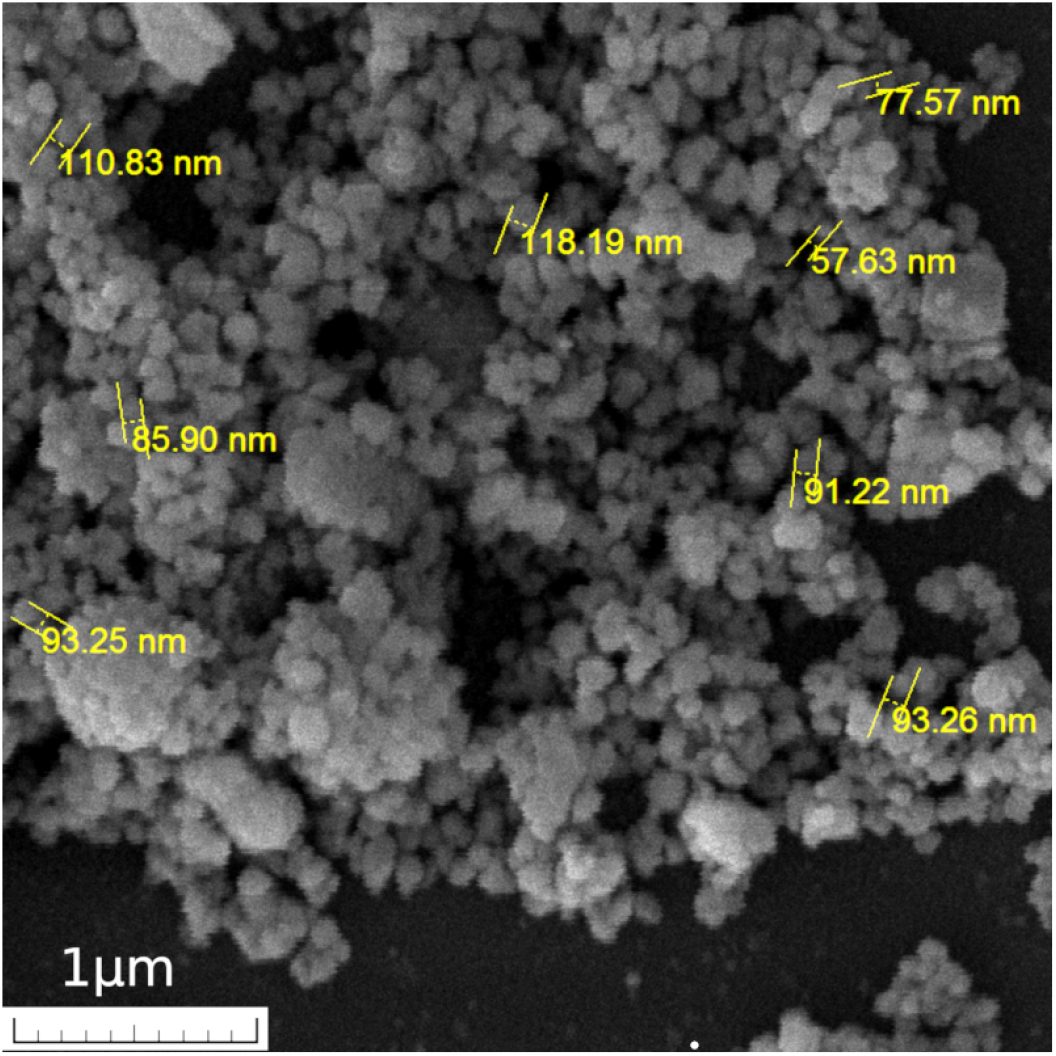
Scanning Electron Microscope images of Zirconia powder obtained by Wet Grinding process (avg. size 90nm, 50,000 x magnification).

### 6.3. Particles’ Distribution in Composite

Figure 4 shows the SEM image of the produced nanocomposite samples. The particles’ distribution in S0, S1, S2 samples containing 0, 3, and 6 phr of Zirconia, respectively, was relatively uniform, and no evidence of particle agglomeration is observed in the composites. However, in S3, which contains 9 phr of Zirconia, the agglomeration of HA and Zirconia particles can be seen. More diagram presented in the Appendix section (Figure A3).

**Fig. 4.**
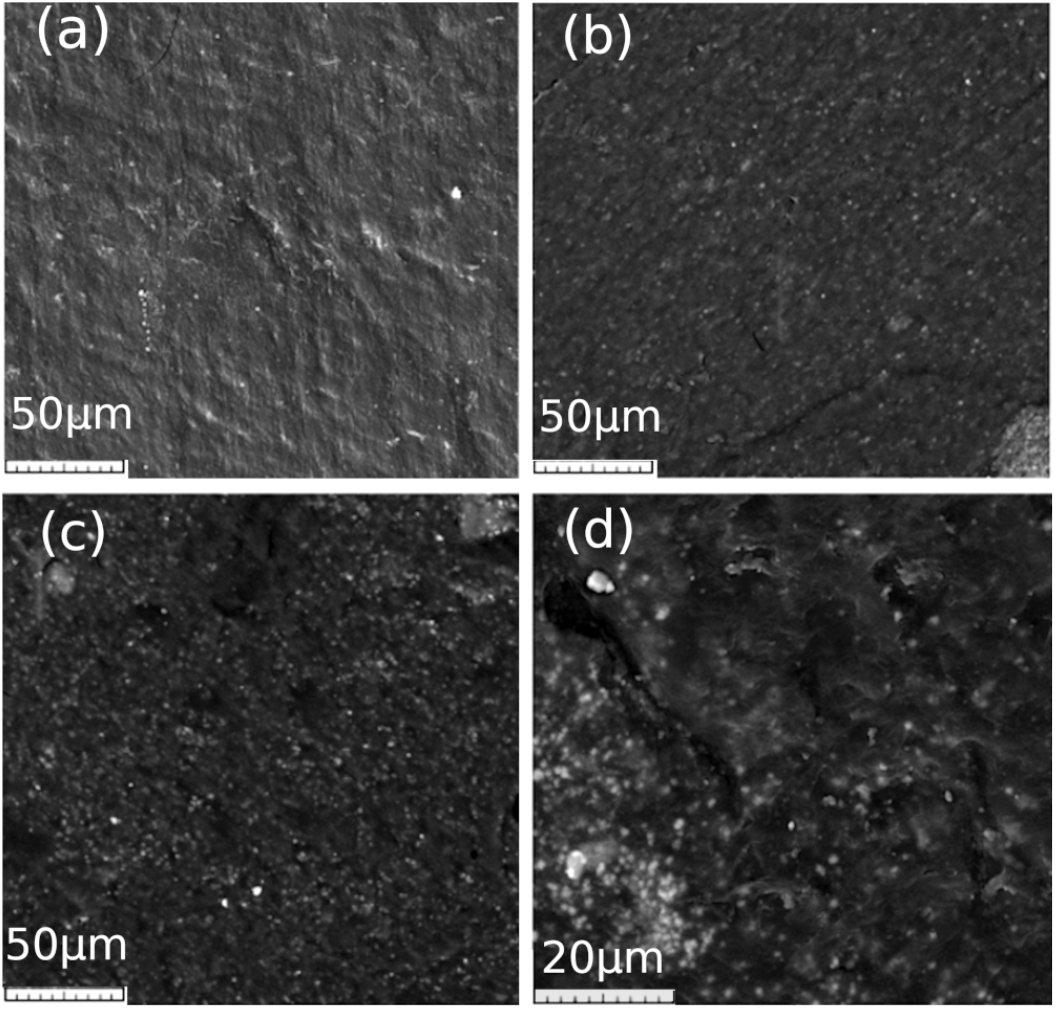
Scanning Electron Microscope images of the produced nanocomposite samples S0 (a), S1 (b), S2 (c), S3 (d) in 1,000 x magnification for S0 to S2 (a, b, c) and 2,900 x magnification for S3 (d).

Fabricated nanocomposites were mapped using the VEGA3 SEM in 5,000x magnification to analyze the reinforcing particles’ distribution more precisely. As shown in Figure 5, samples containing 3 and 6 phr Zirconia (S1 and S2) and samples without Zirconia particles (S0) show relatively good dispersion in the maps but adding 9 phr of Zirconia nano powder leads to a change in sample S3 distribution of additives. The agglomeration of nanopowders can be seen in Figure 5.

**Fig. 5.**
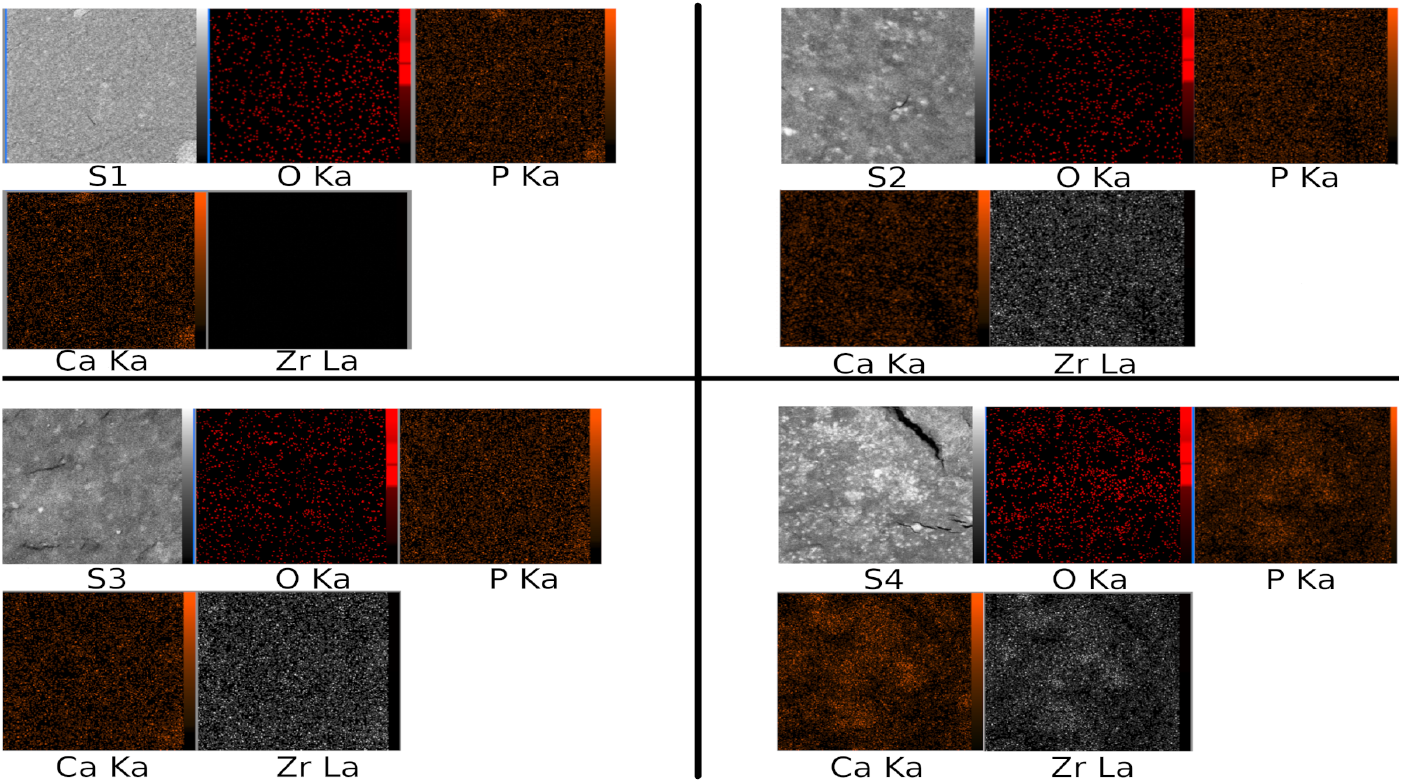
Mapping of four different additives by Scanning Electron Microscopy (5,000 x magnification) in S0 (a), S1 (b), S2 (c), S3 (d).

S3 was scanned by ZEISS DSM-960A FESEM with 21,000 x magnification to see the agglomeration of contents in Figure 6.

**Fig. 6.**
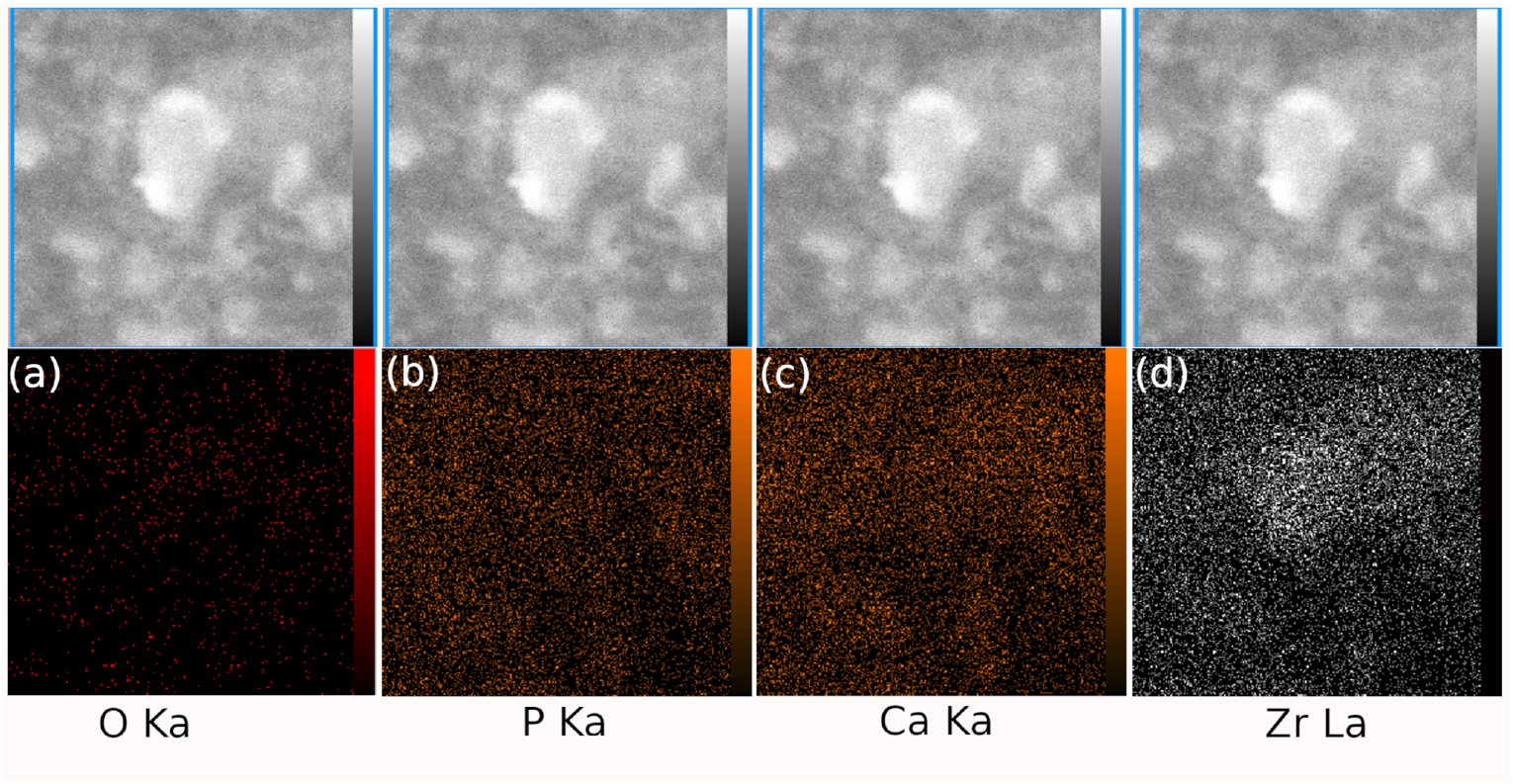
Mapping by Scanning Electron Microscopy (21,000 x magnification) exhibits the distribution of (a) Oxygen, (b) Phosphorous, (c) Calcium, and (d) Zirconium in S3.

As demonstrated in Figure 6, the particle contains a considerable amount of Zirconia and the relative accumulation of HA. The more Zirconia in nanopowder, the more agglomeration occurs due to the intrinsic tendency of nanopowders.

### 6.4. Functionalized nanotubes FTIR

The presence of carboxyl groups can be seen in functionalized carbon nanotubes’ FTIR in Figure 7. Comparing FTIR results of conventional CNT and functionalized CNTs shows the differences between transmission percentage in a wide range of wave numbers representing carboxyl group well-attached to the MWCNTs through the acid treatment.

**Fig. 7.**
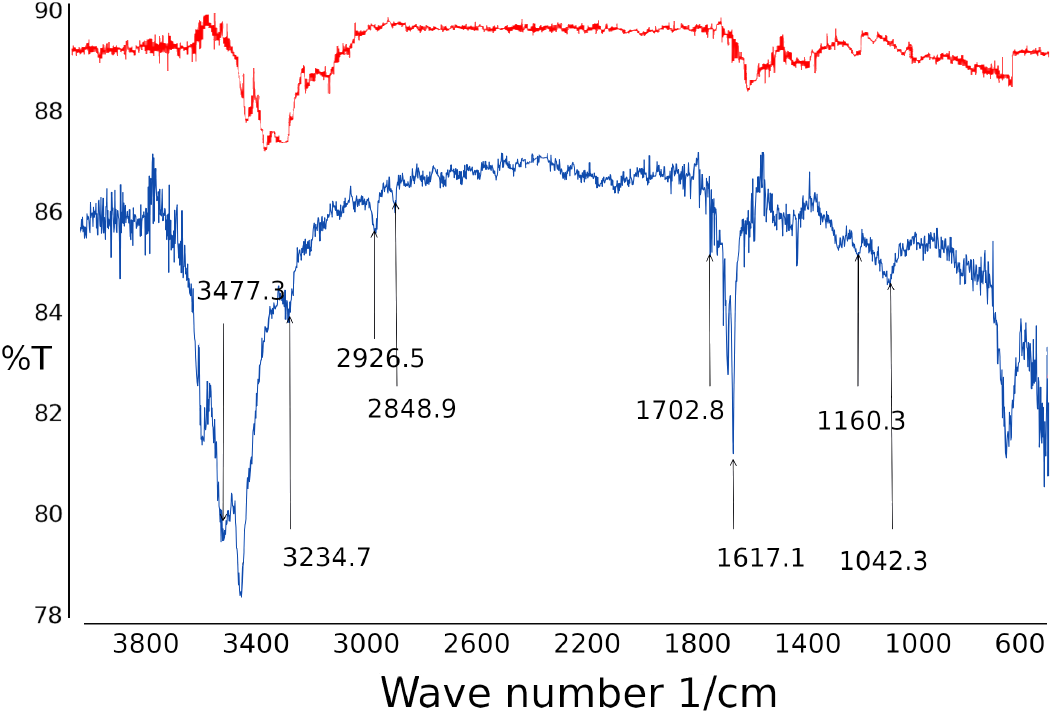
FTIR spectroscopy of conventional carbon nanotubes (red) and functionalized carbon nanotubes (blue).

### 6.5. Nanoindentation Result

The nanoindentation test is one of the most important methods for achieving the hardness and mechanical properties of nano-dimension samples, performed by a very high precision device.

This test provides us with a reduced modulus (representative of the compressive modulus) and the specimens’ hardness.

The reduced modulus and the hardness of the sample are calculated based on two crucial factors: the indenter force and penetration depth at microscales and nanoscales.

By substituting the values in (2), it can be concluded that the achieved (effective) reduced modulus is in a direct relationship with the compressive modulus of our samples.

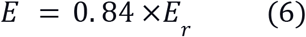

As shown, and concerning the graphs obtained from the nanoindentation test in Figure 8, we can see noticeable changes in the composite compressive modulus and its mechanical properties due to the rise of Zirconia weight percentage.

**Fig. 8.**
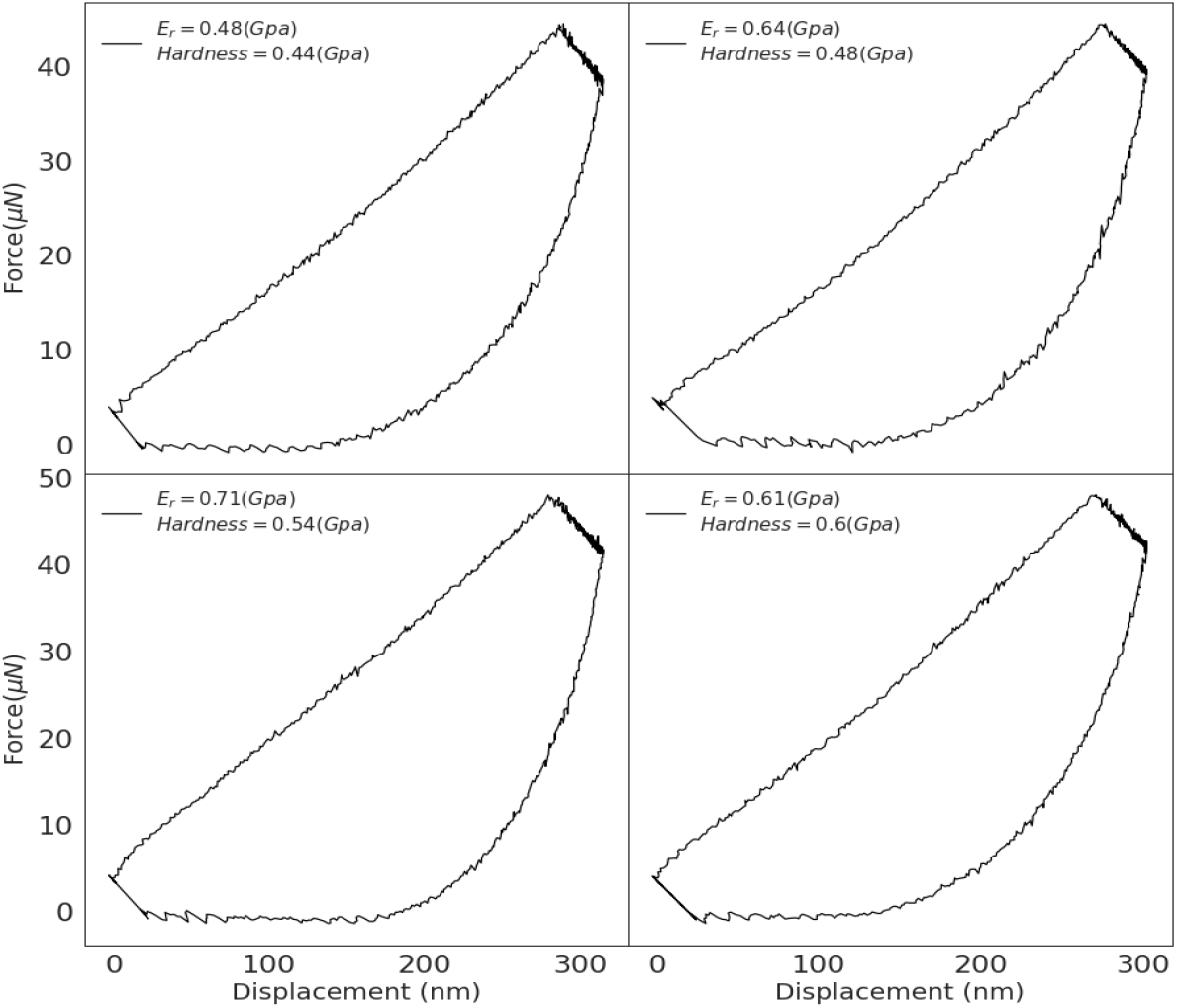
Comparison between reduced modulus and hardness rate in nanoindentation test in (a) S0, (b) S1, (c) S2, and (d) S3. An increase in Zirconia weight percentage positively correlates with the hardness rate, however, the reduced modulus fluctuates between 0.44 (S0) Gpa and 0.6 Gpa (S3).

However, not every elastic modulus follows an upward trend. As we can see in the nanoindentation plot, there is no significant growth in the reduced modulus in S3, but there is a noticeable decrease (14% in comparison with S2) despite a raise of 3 phr of Zirconia.

Also, SEM images of nanocomposites demonstrate the heterogeneous dispersion of HA and Zirconia in S3. So stress concentration and mechanical properties diminishing in these areas could be scientifically feasible.

**TABLE2:** comparison between reduced elastic modulus and hardness obtained from Nanoindentation Test.

| Samples | Hardness<br>(GPa) | $E_r$ (Reduced elastic modulus)<br>(GPa) |
| --- | --- | --- |
| S0 | 0.044 | 0.48 |
| S1 | 0.048 | 0.64 |
| S2 | 0.054 | 0.71 |
| S3 | 0.059 | 0.61 |

Based on the Archard equation and number of reports [**25**], wear rate and hardness are inversely proportional. Therefore, wear rate reduction can be expected by increasing S1, S2, and S3 hardness due to the rise in weight percentage of Zirconia in nanocomposite samples. More diagrams with results are provided in the Appendix section (Figure A1).

### 6.6. Nanoscratch Test

The test results obtained from the Nanoscratch test are based on two important factors, namely force and displacement.

As demonstrated, the nanoscratch diagram (Figure 9) has a horizontal and a vertical axis, that depicts the nano-indenter tip displacement and the lateral to vertical force ratio recorded by the device, respectively. The friction coefficient can be calculated by dividing the vertical force by the lateral force. However, this friction coefficient is a proper measure only when the thin layer coating is employed on material surfaces, or when the loading is negligible.

**Fig. 9.**
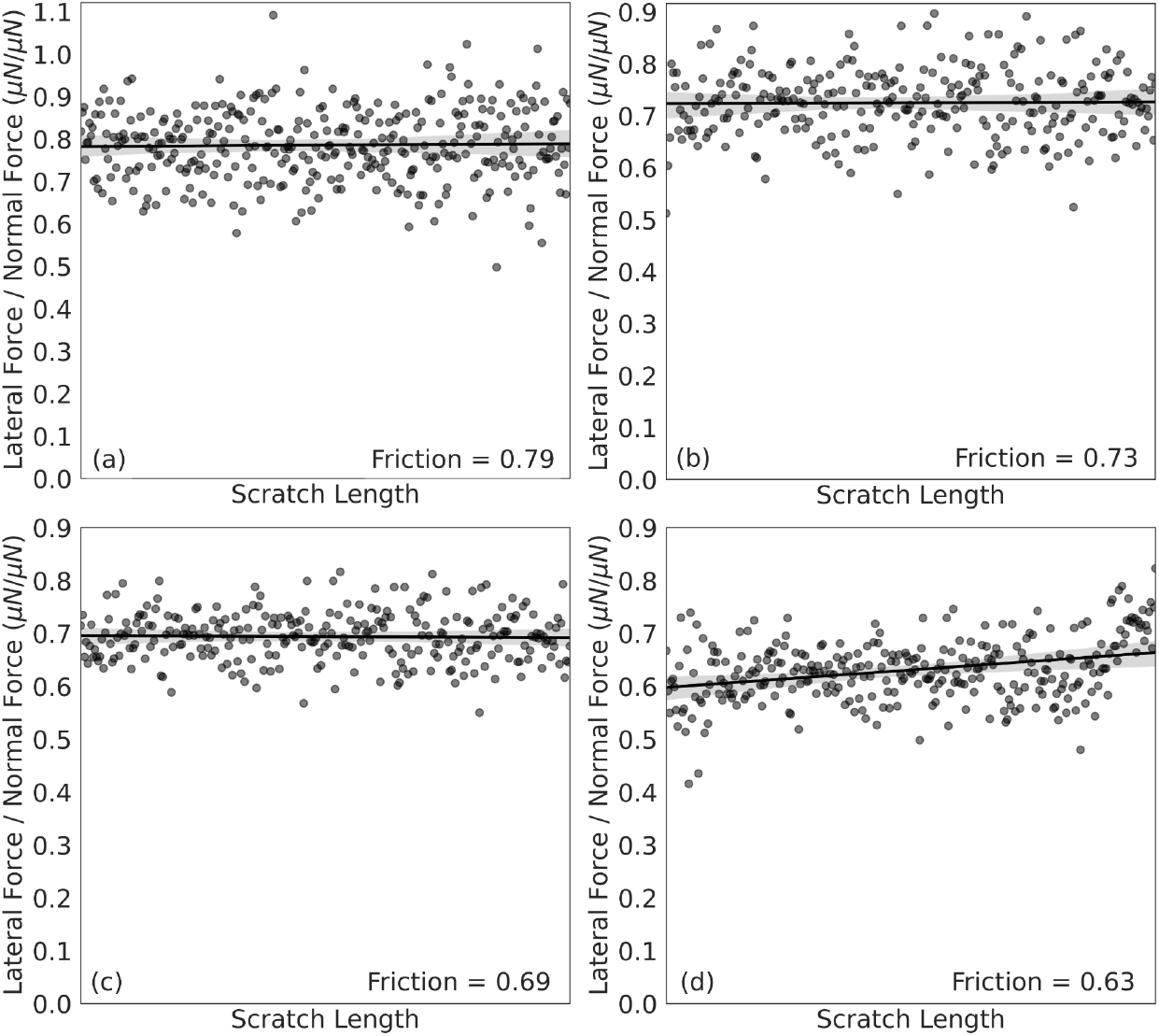
Friction coefficient of five pooled nanoscratch tests: (a) S0, (b) S1, (c) S2, and (d) S3 (d). The horizontal line represents the average of lateral/normal force ratio. The grey shades represent the standard error values.

The friction coefficient and hardness are two significant criteria for wear resistance [**21**,**22**]. According to the graphs, the friction coefficient decreased by increasing Zirconia content in the composite samples from 0.79 phr to 0.63 phr.

This decrease in friction coefficient can be attributed to the composite’s integrity and the significant interface strength of additives within the matrix. As far as we know, the only difference between the four samples is the amount of ZrO_2_ nanopowder that we added to the nanocomposite. So it can be concluded that adding Zirconia as an additive can improve the tribological properties such as wear resistance, hardness, and mechanical properties.

### 6.7. Plasticity index

The plasticity index (PI) represents the material’s elastic recovery ability after removing the deformative load. The value of plasticity index (ψ) for viscoelastic-plastic materials such as polymers fluctuates between 0 and 1, while PI = 1 for fully plastic and PI = 0 for fully elastic materials. Plasticity depends on two key factors such as surface roughness and hardness. Materials with higher ψ have higher friction coefficient and higher friction wear volume.

As shown in Figure 10, by increasing the Zirconia parts per hundred resin/rubber, PI is decreasing. Compared to the previous reported results, relative PI fall can be seen changing from almost 0.6 to 0.45, which is governed by adding MWCNTs and Zirconia nano powder as two other additives [**25**]. It can be due to enhancing the matrix/reinforcements interface due to the nanocomposite fabrication process (Melt Mixing). Besides, the role of Zirconia and MWCNTs in decreasing the PI should not be neglected.

**Fig. 10.**
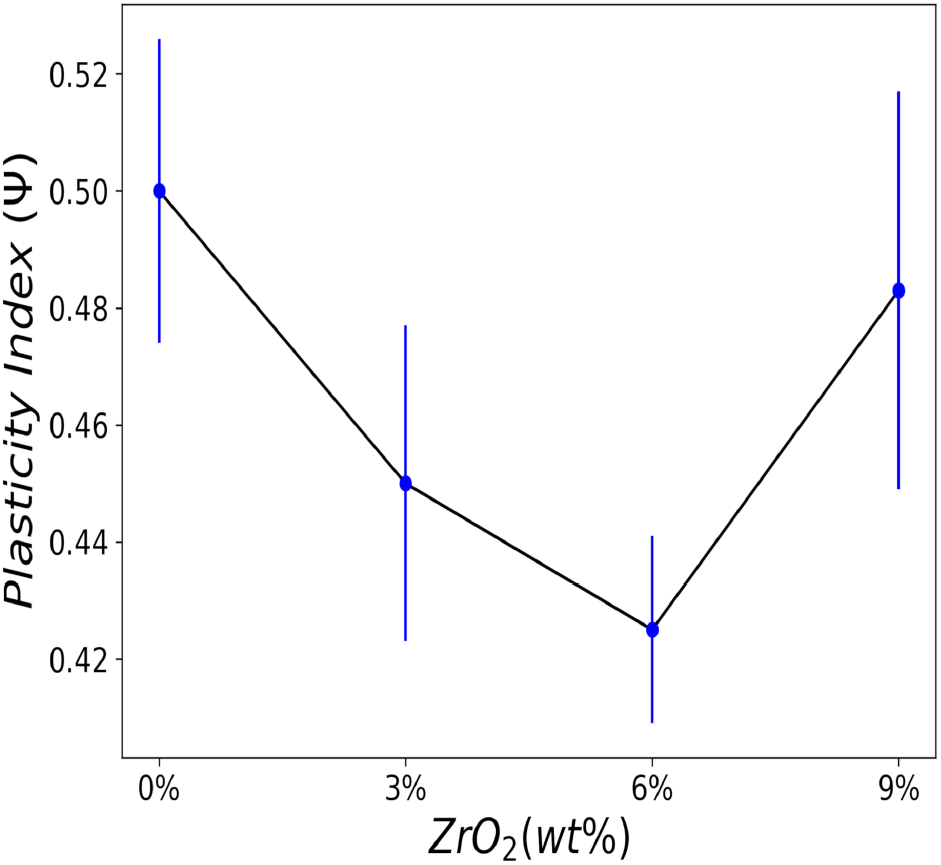
Calculated PI of S0-S3 which includes 0, 3, 6 and 9 phr of Zirconia.

### 6.8. Cytotoxicity Tests (Biological Properties)

Figure 11 shows the MTT assay results for nanocomposite samples compared to the control sample. Clearly, not only cells have high viability in the vicinity of all the samples, but the cell viability rate increased in S0, S1, and S2.

**Fig. 11.**
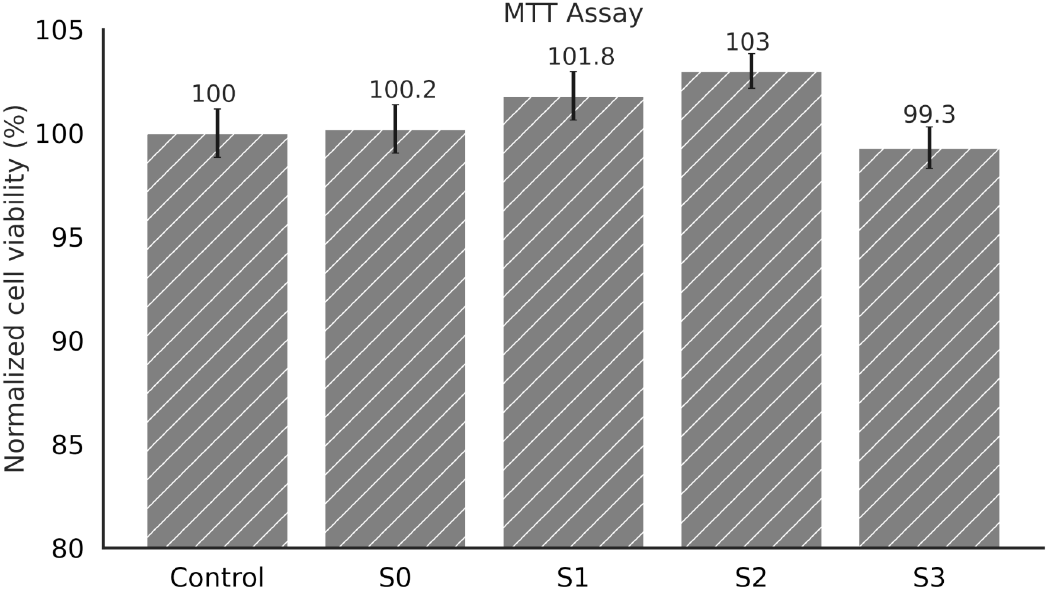
MTT assay of MG-63 cell line in medium exposed to S0, S1, S2, S3, compared to the control (MG-63 cell culture without any composite exposure). The MTT assay was performed on three biological replicates (each with three technical replicates) during three consecutive weeks.

As we know, Zirconia is widely used as a bioinert ceramic in medical applications. Similarly, other composite components, as reported by other researchers, have no detrimental effect on the samples’ biocompatibility [**26**].

### 6.9. Cell Adhesion

In addition to the MTT assay, the cell adhesion test was chosen to assure us about biocompatibility.

As shown in Figure 12, the cells are anchored on the nanocomposite (containing all the composite components) sample, which is considered phase 2 and 3 of the cell adhesion process. Besides, all parts of the nanocomposite are reported to be biocompatible. These two tests, however, are not enough to prove that our fabricated nanocomposites are biocompatible. It needs further investigation.

**Fig. 12.**
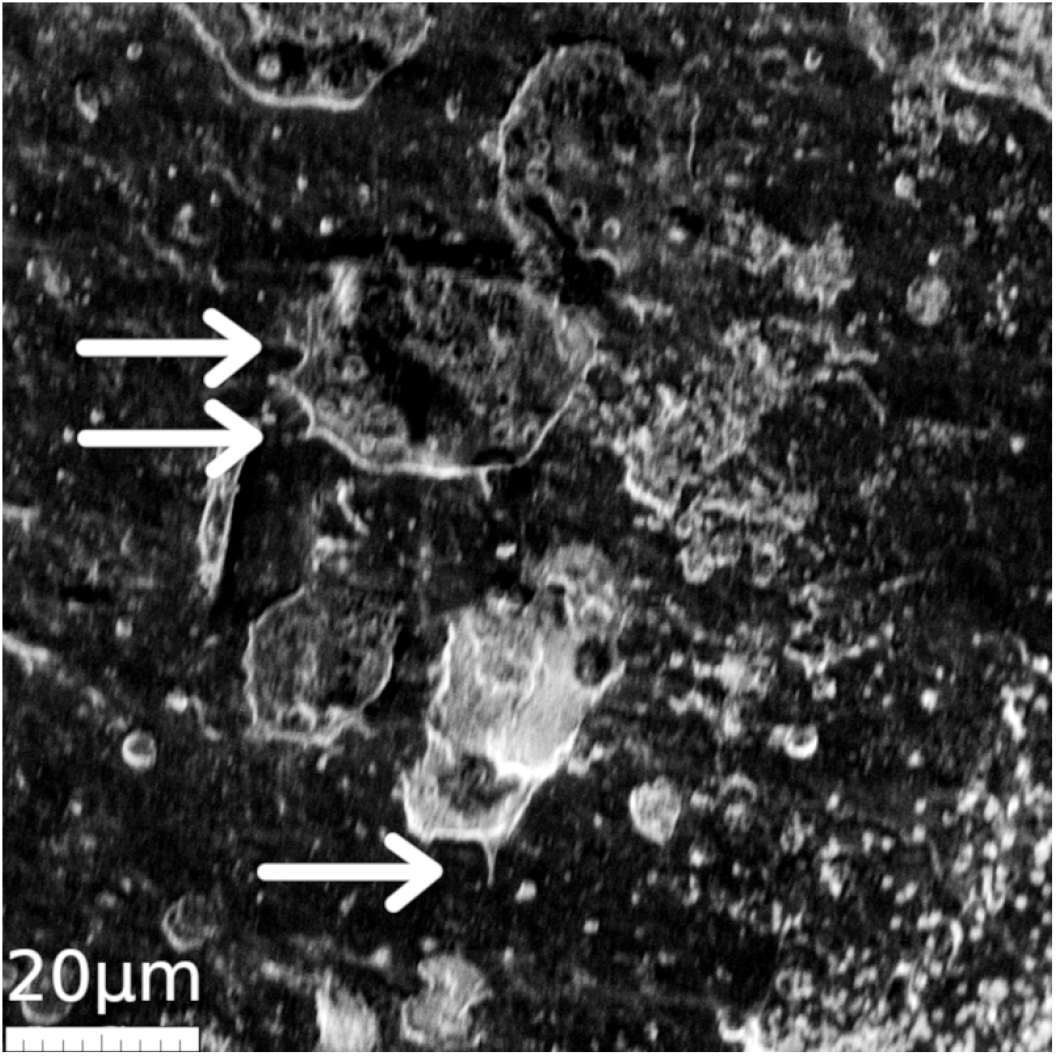
Scanning Electron Microscope image of cell expansion on S3 surface.

## 7. Conclusion

Considering the above-mentioned discussion, it can be concluded that:

First, according to the results drawn from Nanoindentation and Nanoscratch test, increasing Zirconia content results in a rise in hardness and a reduction in friction coefficient in all samples. Secondly, despite the homogeneous distribution in S1 and S2 (containing 3 phr and 6 phr of Zirconia), the agglomeration phenomenon was observed in S3 (with 9 phr). Therefore, the optimum amount of Zirconia nanopowder should be considered to prevent agglomeration, stress concentration, and a drop in mechanical properties, such as compressive elastic modulus.

Thirdly, although the melt mixing method is a time-consuming and costly procedure to fabricate hybrid nanocomposite, the particles’ homogeneous distribution illustrates the method’s advantages in hybrid composite fabrication.

Finally, as discussed above, all samples showed clear responses to the MTT assay and Cell Adhesion Test, yet from the biocompatibility viewpoint, further investigations are needed to prove nanocomposites’ biocompatibility notwithstanding.

## Acknowledgments

We are also immensely grateful to Dr. Setayesh Yasami-khiabani for their comments on an earlier version of the manuscript, although any errors are our own and should not tarnish the reputations of these esteemed persons.

## REFERENCES

[1] S. Sivananthan, S. Goodman, and M. Burke, “11 - Failure mechanisms in joint replacement* A2 - Revell, P.A,” in Joint Replacement Technology (Second Edition), Woodhead Publishing, 2014, pp. 370–400.

[2] S. Kurtz, K. Ong, E. Lau, F. Mowat, and M. Halpern, “Projections of primary and revision hip and knee arthroplasty in the United States from 2005 to 2030,” J Bone Jt. Surg Am, vol. 89, no. 4, pp. 780–785, 2007.

[3] J. Jac Faripour Maybody, A. Nemati, E. Salahi, “Synthesis and properties of MWCNT-HAP composites via SOL-GEL technique” Iranian Journal of Materials Science and Engineering, vol. 10, 2013

[4] J..B. Park, J.D. Bronzino, Biomaterials principles and applications. CRC Press, 1^st^ ed., 2002.

[5] S. M. Kurtz, “The Origins of UHMWPE in Total Hip Arthroplasty,” pp. 33–44, 2016.

[6] S. Teramura, H. Sacoda, T. Terao, M. M. Endo, K. Fujiwara, N. Tomita, “Reduction of Wear Volume from Ultrahigh Molecular Weight Polyethylene Knee Components by the Addition of Vitamin E,” Orthop Res, 2008 Apr;26(4):460–4. doi: 10.1002/jor.20514..

[7] A. A. Edidin and S. M. Kurtz, “Influence of mechanical behavior on the wear of 4 clinically relevant polymeric biomaterials in a hip simulator,” J. Arthroplasty, vol. 15, no. 3, pp. 321–331, Apr. 2000.

[8] P. Steven, M. Kurtz, “From Ethylene Gas to UHMWPE Component: The Process of Producing Orthopedic Implants,” in UHMWPE Biomaterials Handbook 3rd Edition, 2016.

[9] J. A. Puértolas and S. M. Kurtz, “21 - UHMWPE Matrix Composites,” in UHMWPE Biomaterials Handbook (Third Edition), Oxford: William Andrew Publishing, 2016, pp. 369–397.

[10] W. J. Wood, R. G. Maguire, and W. H. Zhong, “Improved wear and mechanical properties of UHMWPE–carbon nanofiber composites through an optimized paraffin-assisted melt-mixing process,” Compos. Part B Eng., vol. 42, no. 3, pp. 584–591, 2011.

[11] Y.-S. Zoo, J.-W. An, D.-P. Lim, and D.-S. Lim, “Effect of carbon nanotube addition on tribological behavior of UHMWPE,” Tribol. Lett., vol. 16, no. 4, pp. 305–309, 2004.

[12] F. S. Senatov, MV. Gorshenkov, SD Kaloshkin, “Biocompatible polymer composites based on ultrahigh molecular weight polyethylene perspective for cartilage defects replacement,” J. Alloys Compd., vol. 586, Suppl, pp. S544–S547, 2014.

[13] S. Kwak et al., “Effect of γ-Ray irradiation on surface oxidation of ultra-high molecular weight polyethylene/zirconia composite prepared by in situ Ziegler-Natta polymerization,” Macromol. Res., vol. 17, no. 8, pp. 603–608, 2009.

[14] A. Gupta, G. Tripathi, D. Lahiri, and K. Balani, “Compression Molded Ultra High Molecular Weight Polyethylene–Hydroxyapatite–Aluminum Oxide–Carbon Nanotube Hybrid Composites for Hard Tissue Replacement,” J. Mater. Sci. Technol., vol. 29, no. 6, pp. 514–522, 2013.

[15] M. Salari, S.M. Taromsari, R. Bagheri and M. A. F. Sani, “Improved wear, mechanical, and biological behavior of UHMWPE-HAp-zirconia hybrid nanocomposites with a prospective application in total hip joint replacement” J. of Materials Science, vol 54, pp 4259–4276, 2019

[16] H. Park, S. Kwak, and S. Kwak, “Wear-Resistant Ultra High Molecular Weight Polyethylene/Zirconia Composites Prepared by in situ Ziegler-Natta Polymerization,” Macromol. Chem. Phys., vol. 206, no. 9, pp. 945–950, 2005.

[17] R. D. A. Maya, J. John and S. Thomas, “Natural Fibre Reinforced Polymer Composites from Macro to Nanoscale - Chapter12: Hybrid composites,” Researchgate-Old city publishing, 2009.

[18] M. Sattari, M. R. Naimi-Jamal, and A. Khavandi, “Interphase evaluation and nano-mechanical responses of UHMWPE/SCF/nano-SiO2 hybrid composites”, Polym. Test., vol. 38, pp. 26–34, 2014.

[19] S. Jain, “Processing of hydroxyapatite by biomimetic process” Ph.D. Thesis, Department of Ceramic Engineering, National Institute of Technology of Rourkela, 2010

[20] N. Kotake, M. Kuboki, S. Kiya, and Y. Kanda, “Influence of dry and wet grinding conditions on fineness and shape of particle size distribution of product in a ball mill,” Adv. Powder Technol., vol. 22, no. 1, pp. 86–92, Jan. 2011.

[21] W. C. Oliver and G. M. Pharr, “An improved technique for determining hardness and elastic modulus using load and displacement sensing indentation experiments,” J. Mater. Res., vol. 7, no. 6, pp. 1564–1583, 1992.

[22] S. A. Mirsalehi, A. Khavandi, S. Mirdamadi, M. R. Naimi-Jamal, and S. M. Kalantari, “Nanomechanical and tribological behavior of hydroxyapatite reinforced ultrahigh molecular weight polyethylene nanocomposites for biomedical applications,” J. Appl. Polym. Sci., vol. 132, no. 23, pp. 1–11, 2015

[23] Lu, Xianjiu Meng, Qingen Wang, Jing Jin, Zhongmin “Transient viscoelastic lubrication analyses of UHMWPE hip replacements” Tribology International., vol 128, pp. 271–278, 2018.

[24] K He, N Chen, C Wang L Wei, J Chen, “Method for determining crystal grain size by x-ray diffraction” Tribology International., vol 103, pp 620–628, 2016.

[25] S. Mosleh-Shirazi, F. Akhlaghi, D.Y. Li, “Effect of graphite content on the wear behavior of Al/2SiC/Gr hybrid nano-composites respectively in the ambient environment and an acidic solution” Tribology International., vol 103, pp 620–628, 2016.

[26] S. Affatato, A. Ruggiero, and M. Merola, “Advanced biomaterials in hip joint arthroplasty. A review on polymer and ceramics composites as alternative bearings,” Compos. Part B Eng., vol. 83, pp. 276–283, 2015

