## supplemental figures for "Fabrication and Evaluation of Mechanical and Tribological Properties of ultra-high-molecular-weight polyethylene-based Nanocomposites Reinforced with Hydroxyapatite, Zirconia and Multi-walled Carbon Nanotubes"

### Appendix:

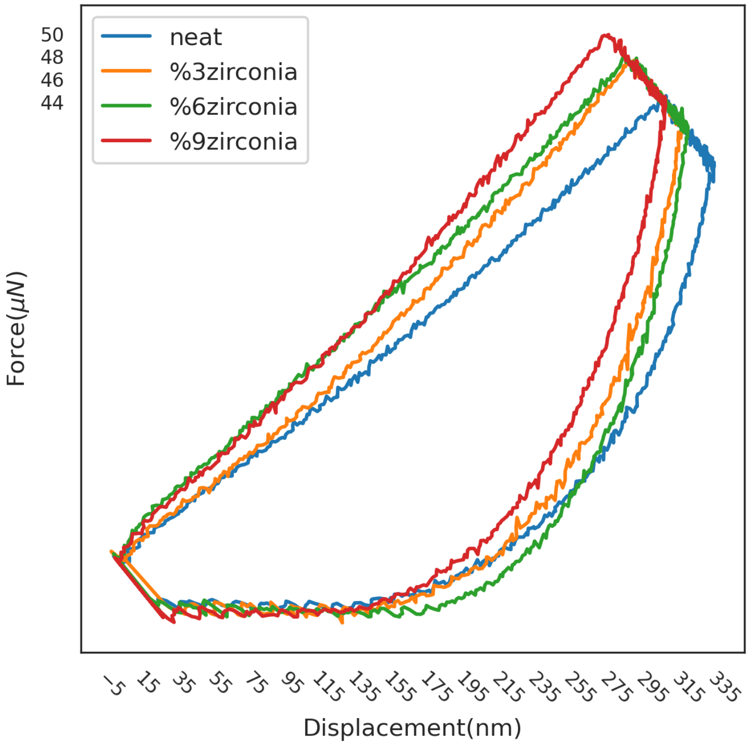


Fig. A1. Comparison between all samples based on reduced modulus and hardness rate in nanoindentation test.

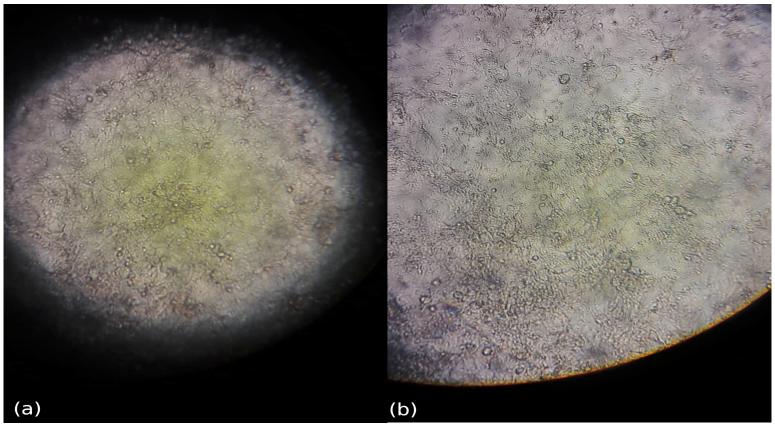


Fig. A2. The MG-63 cell morphology, after (a) 24 hours culture with no extracted sample and (b) exposure to the extracted S3 for 24 hours.

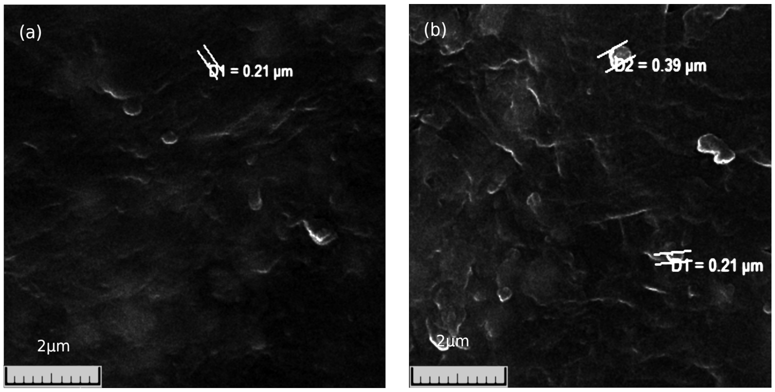


Fig. A3. Particle sizes in neat and 3 wt% composites: (a) HA particle size in S0 (b), HA particle size in S1 (bottom), Zirconia particle size (top)
